# Stimulus-independent neural coding of event semantics: Evidence from cross-sentence fMRI decoding

**DOI:** 10.1101/2020.10.06.327817

**Authors:** Aliff Asyraff, Rafael Lemarchand, Andres Tamm, Paul Hoffman

**Author notes:** These authors contributed equally to the study and are joint first authors (Aliff Asyraff, Rafael Lemarchand). Correspondence to: Dr. Paul Hoffman, School of Philosophy, Psychology & Language Sciences, University of Edinburgh, 7 George Square, Edinburgh, EH8 9JZ, UK.

## Abstract

Multivariate neuroimaging studies indicate that the brain represents word and object concepts in a format that readily generalises across stimuli. Here we investigated whether this was true for neural representations of simple events described using sentences. Participants viewed sentences describing four events in different ways. Multivariate classifiers were trained to discriminate the four events using a subset of sentences, allowing us to test generalisation to novel sentences. We found that neural patterns in a left-lateralised network of frontal, temporal and parietal regions discriminated events in a way that generalised successfully over changes in the syntactic and lexical properties of the sentences used to describe them. In contrast, decoding in visual areas was sentence-specific and failed to generalise to novel sentences. In the reverse analysis, we tested for decoding of syntactic and lexical structure, independent of the event being described. Regions displaying this coding were limited and largely fell outside the canonical semantic network. Our results indicate that a distributed neural network represents the meaning of event sentences in a way that is robust to changes in their structure and form. They suggest that the semantic system disregards the surface properties of stimuli in order to represent their underlying conceptual significance.

## Introduction

Most neuroscientific theories of semantic representation hold that meanings are coded, at least in part, independently of the stimuli used to elicit them (Binder & Desai, 2011; Meteyard, Cuadrado, Bahrami, & Vigliocco, 2012; Patterson, Nestor, & Rogers, 2007; Rogers et al., 2004; Simmons & Barsalou, 2003). These theories propose, for example, that the same semantic representation for the concept *DOG* is engaged whether one reads the word “dog”, sees a canine in the park or hears the sound of barking. This position is most strongly associated with hub-and-spoke theories (Hoffman, McClelland, & Lambon Ralph, 2018; Lambon Ralph, Jefferies, Patterson, & Rogers, 2017; Rogers et al., 2004). These emphasise the role of supramodal semantic representations that abstract away from perceptual inputs in order to code deeper conceptual structure. Embodied approaches to semantics place greater emphasis on the role of sensory-motor simulations in comprehension (Barsalou, 1999; Pulvermüller, 2013; Zwaan, 2004), which are linked more directly with specific modalities (e.g., engagement of the motor system when people comprehend action words; Hauk, Johnsrude, & Pulvermuller, 2004). This view is also compatible with stimulus-independent representation, since the same simulations might be activated by a range of different stimuli. The emergence of multivariate neuroimaging techniques has afforded new opportunities to assess how and where stimulus-independent semantic representations are coded in the brain. In one of the first such fMRI studies, Fairhall and Carramazza (2013) presented participants with pictures of objects belonging to different taxonomic categories and with their written names. They tested for cross-modal semantic representation by training a multivariate classifier to discriminate the categories using neural responses to the pictures and then testing its ability to classify the category of the word stimuli (and vice-versa). Other studies have also searched for commonalities in the neural patterns for objects elicited by word and pictures (Devereux, Clarke, Marouchos, & Tyler, 2013; Shinkareva, Malave, Mason, Mitchell, & Just, 2011), for neural similarities between auditory and written words (Liuzzi et al., 2017) and for convergence in the neural responses to the same concepts presented in two different languages (Correia et al., 2014). In general, these studies converge in identifying a range of semantic processing regions distributed throughout frontal, temporal and parietal cortices, all of which code the underlying conceptual significance of a stimulus, independent of the particular form the stimulus takes. This is in keeping with theoretical approaches holding that conceptual knowledge is coded in one or more semantic hubs, situated downstream from sensory and motor cortices (Binder & Desai, 2011; Lambon Ralph et al., 2017; Margulies et al., 2016).

Although there is evidence for stimulus-independent representation of individual words and objects, fewer studies have investigated whether more complex representations, expressed at the sentence level, are also coded in a stimulus-independent fashion. This is a critical question because the flexible nature of language allows for events with the same conceptual content to be described in very different ways (see Table 1 for examples). Studies have demonstrated that fMRI activation patterns elicited by sentences do contain information about their semantic content. A number of researchers have developed predictive models of sentence-level activation by decomposing sentences into the semantic properties of their constituents and identifying their neural correlates (Anderson et al., 2017; Frankland & Greene, 2020; Just, Wang, & Cherkassky, 2017; Pereira et al., 2018; Wang, Cherkassky, & Just, 2017). Such models can successfully predict activation patterns to new sentences that describe different events. Activation in left temporal and parietal cortices appears to be critical in supporting such predictions.

**Table 1:** Sentences describing one of the four events

|  | Lexical items 1 | Lexical items 2 |
| --- | --- | --- |
| Active form | The student considered the problem. | The pupil pondered the issue. |
| Passive form | The problem was considered by the student. | The issue was pondered by the pupil. |

Cross-language studies have begun to use similar techniques to investigate whether neural representations of sentence meaning generalise across changes in linguistic form. Hu et al. (2019) trained a multivariate classifier to discriminate between coherent and incoherent sentence pairs presented to bilinguals in Chinese or Japanese script. Patterns in left parietal cortex and inferior prefrontal cortex showed successful cross-language classification, suggesting that a similar neural representation was engaged by both languages. However, in this study the classifier was not required to discriminate the content of the sentences, only whether they described a coherent event. In contrast, Yang et al. (2017) developed a feature-based semantic model to code the content of 60 distinct English sentences. This model was trained with fMRI data from English speakers to form a predictive model of sentence activation patterns across the whole brain. Once trained, the model could successfully predict neural responses to Portuguese translations of the sentences in a new group of Portuguese speakers. This study demonstrated that the areas of the brain activated by particular events are broadly similar even across different individuals who speak different languages. However, it was not designed to identify which specific brain regions demonstrate this stimulus-independent coding. In the present study we aimed to do this in a multivariate fMRI study that held constant the language used to describe events, but systematically varied the linguistic forms of the descriptions.

We investigated neural representations of four simple events, systematically varying the syntactic structures and lexical items used to describe each event. We aimed to identify brain regions whose activity patterns discriminated between different events and then to assess whether this discrimination was robust to changes in the syntactic and lexical properties of the sentences. We predicted that the regions of the left-lateralised canonical semantic network, previously linked with supramodal representation of words and objects, would show stimulus-independent coding of event meanings. We then reversed our analysis, testing for brain regions that could successfully discriminate between syntactic structures, while generalising over the semantics of the events. Finally, we tested for regions that discriminated between different lexical items used to describe the same events. Together, these analysis allowed to distinguish between brain regions that process the conceptual content of sentences and those involved in processing their form.

## Method

### Participants

26 native English speakers took part in the study (20 female; mean age 22.48, range 18-35 years). All were classified as right-handed using the Edinburgh Handedness Inventory (Oldfield, 1971) and none reported dyslexia or any history of neurological illness. All provided written informed consent and the study was approved by University of Edinburgh School of Philosophy, Psychology & Language Sciences Research Ethics Committee.

### Stimuli

We probed neural representations of four coherent events designed to cover a broad range of conceptual knowledge, including living and non-living agents (cow, student; computer, lorry), motion and static verbs (driving, jumping; processing, considering) and concrete and abstract patients (fence, bridge; files, problems). Each event was presented to participants in four different sentence forms that varied in their syntactic structure and lexical items (see Table 1 for examples). Syntactic structure was varied by describing the event with either an active or passive sentence. Lexical items were varied by substituting semantically-similar words to describe each agent, patient and action. Thus, in total there were 16 event sentences used in our multivariate analyses. We also created 16 anomalous sentences that used the same lexical items and syntactic structures as the event sentences but did not describe a coherent event. These were created by reshuffling the constituents of the event sentences (e.g., “the document was considered by the lorry”). To validate this manipulation, 18 participants, who did not take part in the main study, were asked to rate the meaningfulness of each sentence on a five-point scale. Event sentences received significantly higher ratings than anomalous sentences (Meaningful *M* = 4.56, *SD* = 0.32; Anomalous *M* = 1.53, *SD* = 0.57; *t*(30) = 18.6, *p* < 0.001). A full list of the sentences used is provided in Supplementary Table 1.

### Procedure

Participants completed six runs of scanning. In each run, they were presented with each of the 32 sentences once and were asked to decide whether the sentence was meaningful or not. Manual responses were made using the left and right hands, with the mapping of these to response options counterbalanced over participants. Each trial began with a fixation cross presented for 500ms. This was followed by the written sentence, presented in the centre of the screen for 4s. Trials were separated by a jittered inter-stimulus interval of between 4s and 8s (mean 6s). The order of the sentences was fully randomised for each run and for each participant.

### Image acquisition and preprocessing

Images were acquired on a 3T Siemens Prisma scanner using a 32-channel head coil. We employed a whole-brain multi-echo acquisition protocol, in which data was simultaneously acquired at three echo times (13ms, 31ms and 48ms). Data from the three echo series were weighted and combined and the resulting time-series denoised using independent components analysis (ICA). This protocol reduces the influence of motion and other artefacts and, importantly, improves signal quality in regions that typically suffer from susceptibility artefacts, such as the ventral anterior temporal lobes (Kundu et al., 2017). The TR was 1.7s and images consisted of 46 slices with an 80 × 80 matrix and isotropic voxel size of 3mm. Multiband acceleration with a factor of 2 was used and the flip angle was 73°. Six runs of 195 volumes (331.5s) were acquired. A high-resolution T1-weighted structural image was also acquired for each participant using an MP-RAGE sequence with 0.8mm isotropic voxels, TR = 2.62s, TE = 4.5ms.

Images were preprocessed using the TE-Dependent Analysis Toolbox 0.0.7 (Tedana) (DuPre et al., 2019) and SPM12. Motion parameters were estimated using the first echo series, prior to slice-timing correction (as recommended by Power, Plitt, Kundu, Bandettini, & Martin, 2017). Slice-timing correction was then applied, images were interpolated to the space of the first volume using the previously obtained motion estimates, and Tedana was used to optimally combine the three echo series into a single time-series. Tedana’s denoising algorithms were also applied to the data, independently for each scanning run. These use ICA to partition the data and then to classify each component as either BOLD-related or noise-related, based on its pattern of signal decay over increasing TEs (Kundu et al., 2017). Components classified as noise were discarded. Following denoising, SPM was used to unwarp images with a B0 fieldmap, coregister them to the anatomical scans and normalise them to MNI space with DARTEL (Ashburner, 2007).

For initial univariate analyses, images were smoothed with a kernel of 8mm FWHM. Data were treated with a high-pass filter with a cut-off of 128s and the six runs were analysed using a single general linear model. For each run, one regressor modelled presentation of event sentences and another presentation of anomalous sentences. Covariates consisted of six motion parameters and their first-order derivatives.

### Multi-voxel pattern analysis

For multivariate analysis, smoothing of 4mm FWHM was applied to the normalised images. Although multivariate analyses are often performed on unsmoothed images, there is evidence that a small amount of smoothing can slightly improve classifier performance (Gardumi et al., 2016; Hendriks, Daniels, Pegado, & Op de Beeck, 2017). Each run was analysed with a separate general linear model which included a separate regressor for each of the 32 sentences. T-maps were generated for each event sentence, which were then submitted to decoding analyses using CoSMoMVPA (Oosterhof, Connolly, & Haxby, 2016). Our main analyses tested for effects across the whole of the cortex using a spherical searchlight with a radius of four voxels. We also tested for decoding effects in five anatomical regions of interest described later.

We performed five different analyses that investigated the neural representation of different aspects of sentence information. In Analysis 1, we simply tested for regions whose activation patterns could discriminate between sentences describing different events. This gave us some initial information about which brain regions were sensitive to differences between sentences. However, no generalisation to novel sentences was required in this analysis so these effects could be stimulus-specific. Our main tests of the experimental hypothesis came in Analyses 2 and 3. Here, we trained the classifier to decode events using one half of the sentences and tested it using the other half. The two sets of sentences differed either lexically (Analysis 2) or syntactically (Analysis 3). These analyses identified brain regions that coded the conceptual content of the events in a stimulus-independent fashion. Finally, Analyses 4 and 5 tested which brain regions could decode surface features of the sentences (lexical and syntactic) that were orthogonal to event-level semantic information. These analyses allowed us to determine the degree to which areas coding the conceptual content of the events were sensitive to other features of the stimuli.

### Analysis 1: Sentence-specific decoding

The first analysis tested the ability of brain regions to discriminate between sentences that describe different events, without requiring the classifier to generalise over different descriptions. We selected a set of four sentences that described the four events (with the constraint that they all had the same syntactic structure). We trained the model to discriminate between the four sentences and tested it on new instances of the same sentences (see Figure 1A). This process was repeated for all possible combinations of four sentences representing the four events (32 iterations) and the results averaged to give an overall accuracy map for each participant. Data were partitioned into train and test sets with leave-one-run-out cross-validation: the classifier was trained on data from five runs and tested on the remaining run and this process was repeated until each run had served as the test set.

**Figure 1:**
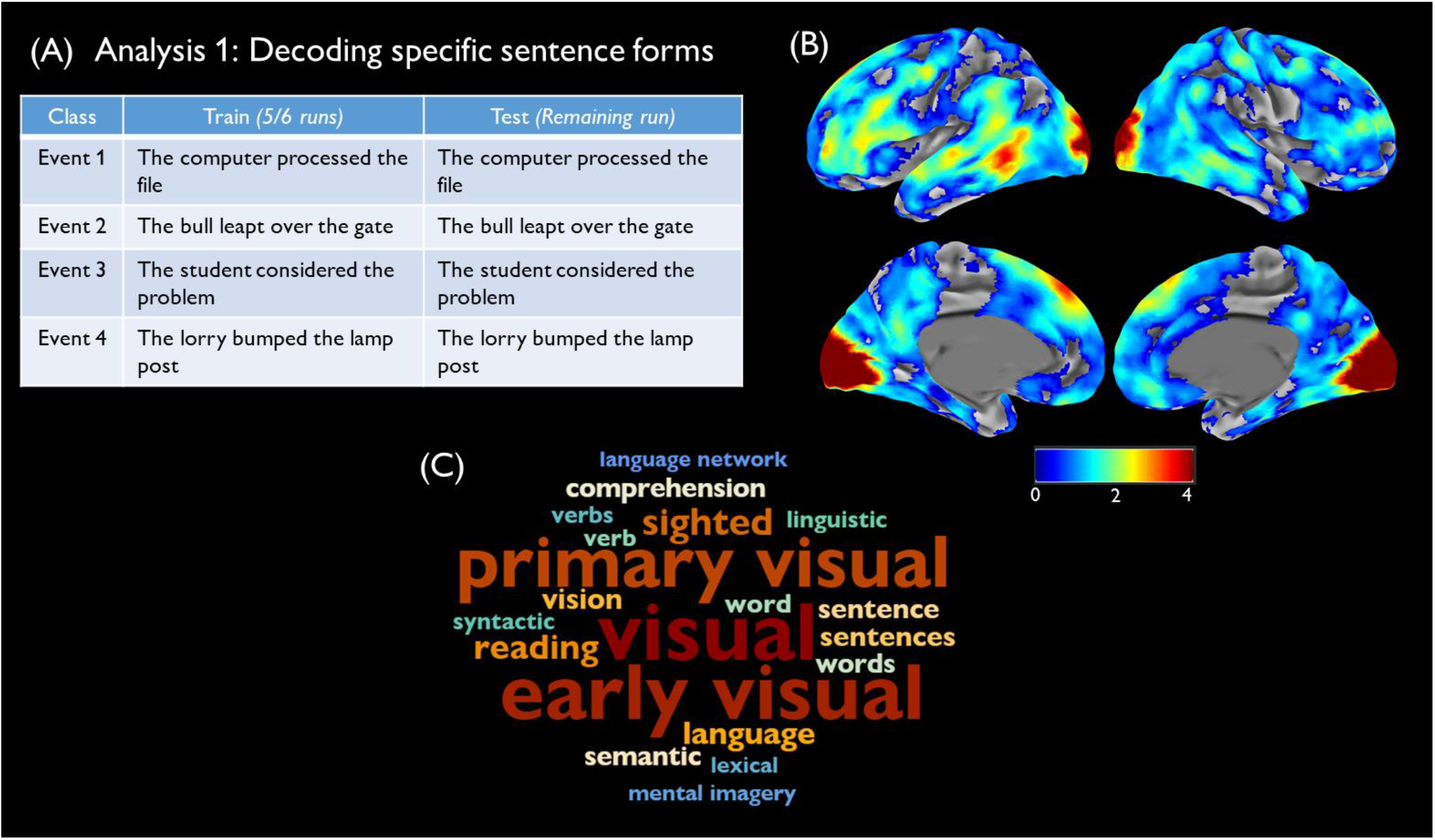
<u>Analysis 1</u> (A) Training and test stimuli for one iteration of the analysis. (B) Decoding accuracy map, relative to chance level. Decoding accuracy is thresholded at cluster-corrected p< 0.05 and maps were smoothed at 5mm FWHM for display purposes. (C) Terms most correlated with accuracy map in Neurosynth.

### Analyses 2 and 3: Generalisation of event identity across sentence forms

The second and third analyses were our primary tests of the experimental hypothesis. Here, we sought brain regions that could decode event information in a way that generalises to new descriptions of the events. Analysis 2 tested generalisation to new lexical items and Analysis 3 generalisation to new syntactic forms. In Analysis 2, the classifier was trained to discriminate the four events, using data from eight sentences. In this set of eight sentences, a single set of lexical items was used to describe each event. The classifier was then tested on the remaining eight sentences, which described the same four events using different lexical items (see Figure 2A). Thus, above-chance decoding could only be achieved where brain regions coded information about the events in a way that generalised to novel sentences. The process was repeated until all possible combinations of lexical items had served as the training set (2^4^ = 16 iterations). Analysis 3 proceeded in a similar fashion, except that train and test sets differed in the syntactic forms used to describe each event (see Figure 2C). Thus, in Analysis 3 the same words were present in train and test sets but they were embedded in different grammatical structures.

**Figure 2:**
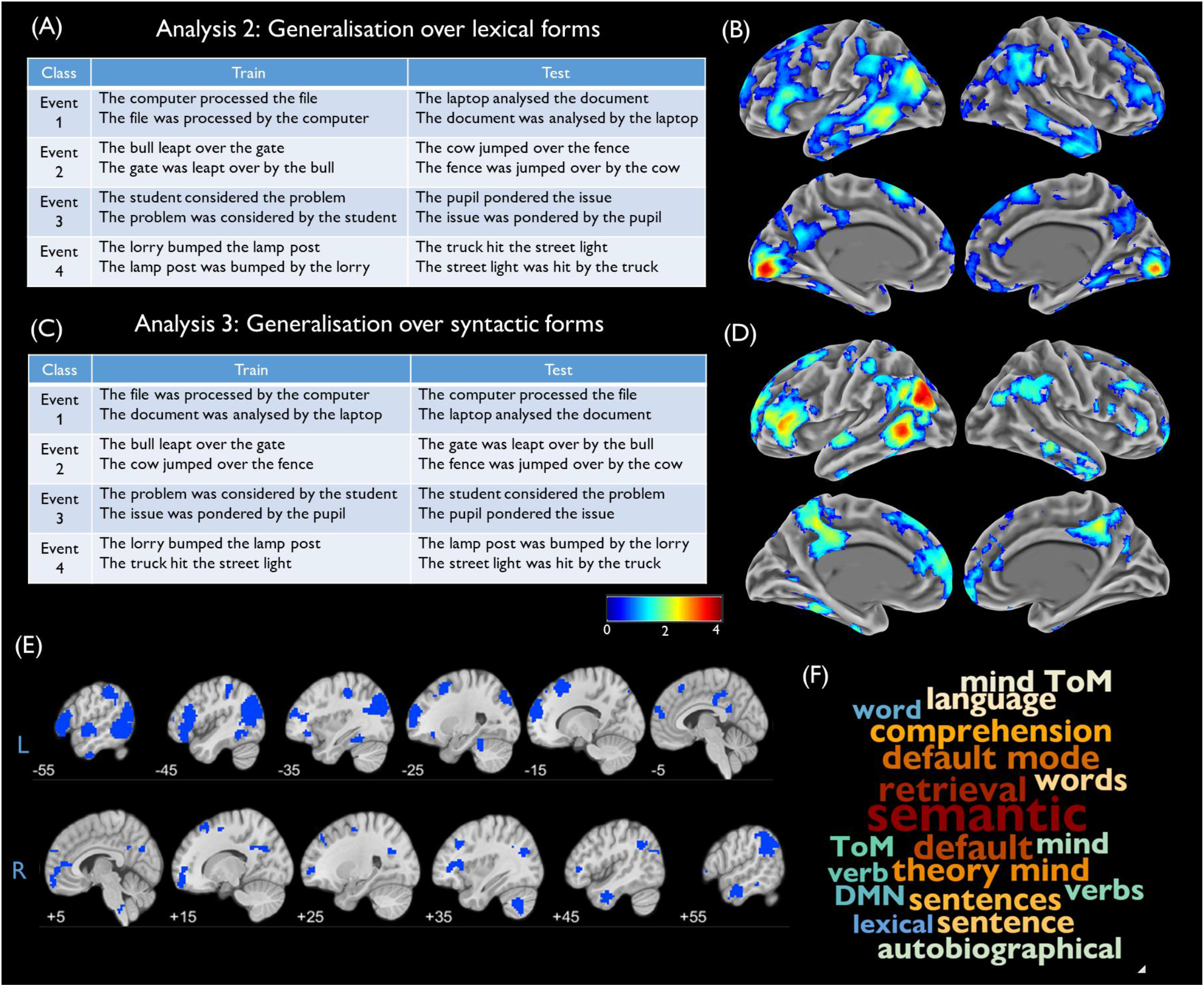
<u>Analyses 2 and 3</u> (A) Training and test stimuli for one iteration of Analysis 2. (B) Decoding accuracy map for Analysis 2, relative to chance level. (C) Training and test stimuli for one iteration of Analysis 3. (D) Decoding accuracy map for Analysis 3, relative to chance level. (E) Conjunction map showing regions that significantly exceeded chance level in both analyses. (F) Terms most correlated with mean accuracy map in Neurosynth. Decoding accuracy is thresholded at cluster-corrected p< 0.05 and maps were smoothed at 5mm FWHM for display purposes.

### Analysis 4: Decoding syntactic structure

Here we reversed the logic of the previous analyses and asked whether any brain regions coded the syntactic structure of the sentences, independent of the events being described. We performed this analysis to explore whether any brain regions coded syntax in a content-independent manner, and whether these regions overlapped with those coding the content of events. The model was trained to discriminate between active and passive sentences. Trials were divided such that eight sentences describing two events were used as the training set and eight sentences describing the remaining two events were used as the test set (see Figure 3A). Thus, above-chance classification could only be achieved if patterns of brain activity discriminated between active and passive sentences in a way that generalised across different events. Decoding was repeated until all possible pairs of events had served as the training set (*C*(4,2) = 6 iterations).

**Figure 3:**
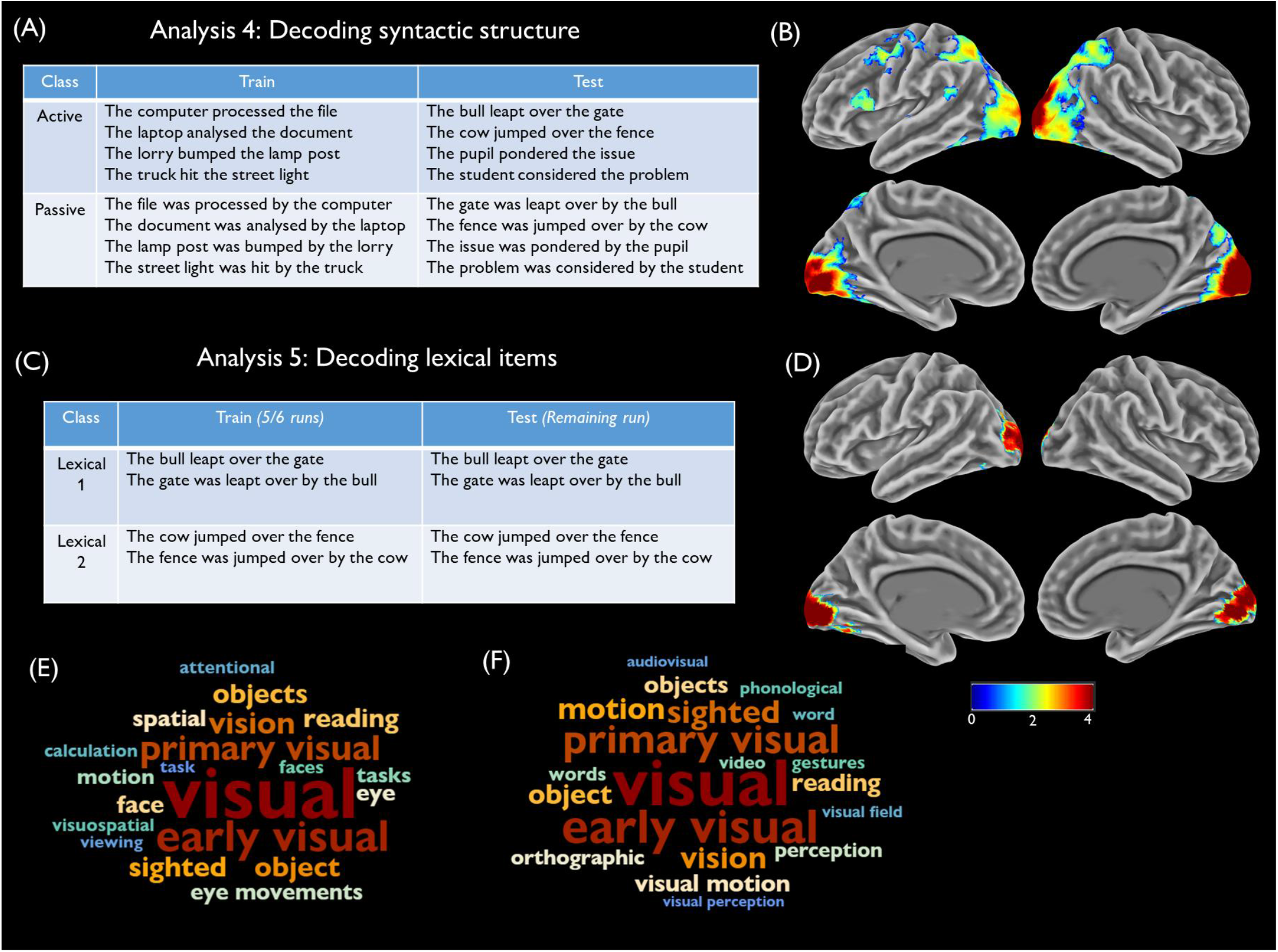
<u>Analyses 4 and 5</u> (A) Training and test stimuli for one iteration of Analysis 4. (B) Decoding accuracy map for Analysis 4, relative to chance level. (C) Training and test stimuli for one iteration of Analysis 5. (D) Decoding accuracy map for Analysis 5, relative to chance level. (E) Terms most correlated with Analysis 4 accuracy map in Neurosynth. (F) Terms most correlated with Analysis 5 accuracy map in Neurosynth. Decoding accuracy is thresholded at cluster-corrected p< 0.05 and maps were smoothed at 5mm FWHM for display purposes

### Analysis 5: Decoding lexical items

Finally, we wanted to test whether any regions coded for the lexical items used in the sentences, independently of the events being described. Here it was not possible to test for generalisation across events as each event was associated with a different set of lexical items. Instead, we treated each event separately and trained the classifier to discriminate between the two sets of lexical items associated with it (see Figure 3C). The classifier was then tested on new instances of the same sentences, with leave-one-run-out cross-validation used to ensure independence of training and testing data. This process was repeated for each of the four events and the results averaged. This analysis tested whether any brain regions discriminated between sentences that differ lexically but convey the same (or very similar) semantic information.

In all analyses, classification was performed using a support vector machine (LIBSVM) with the regularisation parameter *C* set to 1. To determine whether classification was better than expected by chance, permutation tests were performed using the two-stage method introduced by Stelzer et al. (2013). For each participant, we trained and tested the classifier repeatedly on data in which the class labels had been randomly permuted within each run. This process was repeated 100 times (divided equally between all iterations of the training stimuli) to provide an accuracy distribution for each participant under the null hypothesis. Following this, a Monte Carlo approach was taken to generate a null accuracy distribution at the group level. Specifically, from each participant’s null distribution, we randomly selected one accuracy map for each training iteration and averaged these to give a group mean. This process was repeated 10,000 times to generate a distribution of the expected group accuracy under the null hypothesis. In ROI analyses, the position of the observed group accuracy in this null distribution was used to determine a p-value (e.g., if the observed accuracy was greater than 99% of values in the null distribution, this would represent a p-value of 0.01). For searchlight analyses, observed and null accuracy maps were entered into CoSMoMVPA’s Monte Carlo cluster statistics function, which returned a statistical map corrected for multiple comparisons using threshold-free cluster enhancement (Smith & Nichols, 2009). These maps were thresholded at corrected *p* < 0.05.

To aid interpretation of results, unthresholded decoding accuracy maps were submitted to Neurosynth and correlated with its meta-analytic maps (Rubin et al., 2017). Neurosynth is an automated meta-analysis tool that identifies terms commonly used in the neuroimaging literature and relates these to reported activation co-ordinates. A Neurosynth map for a given term indicates the likelihood that activation in each voxel is preferentially associated with studies that use that term (e.g., in the map for “semantic”, each voxel’s value indicates the likelihood that activation is reported there in studies that discuss semantics, relative to studies that don’t). By correlating these maps with our unthresholded decoding maps, we were able to determine which terms are most consistently associated with the set of regions in which we observed strong decoding. To visualise the results, terms relating to anatomical structures were removed and the 20 most correlated terms were extracted and plotted as a word cloud (for similar uses of this technique, see Vatansever et al., 2017; Wang, Margulies, Smallwood, & Jefferies, 2020).

### Regions of interest

ROI analysis focused on left-hemisphere anatomical regions, selected based on their involvement in semantic processing. These are shown in Figure 4A. Four of the five ROIs were defined using probability distribution maps from the Harvard-Oxford brain atlas (Makris et al., 2006), including all voxels with a >30% probability of falling within the following regions:

**Figure 4:**
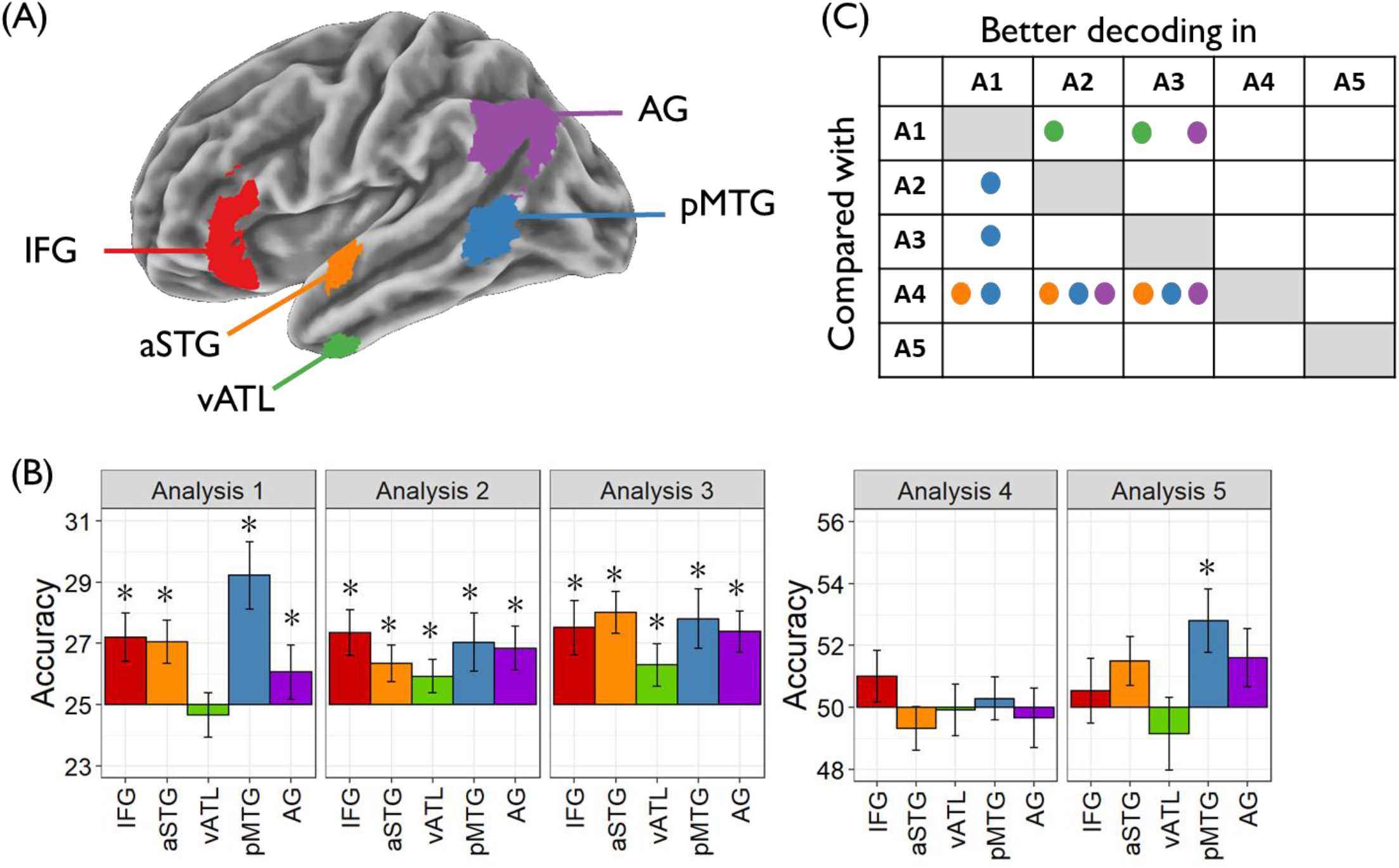
<u>Region of interest analyses</u> (A) Location of anatomical regions of interest. (B) Decoding accuracy for each ROI in each analysis. * indicates accuracy significantly greater than chance (one-tailed p < 0.05, corrected for multiple comparisons using the false discovery rate approach). (C) Results of pairwise comparison of analyses in each ROI. Circles indicate cases where the difference between analyses exceeded that expected by chance (two-tailed p < 0.05, corrected for multiple comparisons using the false discovery rate approach). A1 = Analysis 1 etc.

1. Inferior frontal gyrus (IFG): the pars orbitalis and pars triangularis regions of inferior frontal gyrus, with voxels more medial than x=-30 removed to exclude medial orbitofrontal cortex
2. Anterior superior temporal gyrus (aSTG): the anterior division of the superior temporal gyrus
3. Ventral anterior temporal lobe (vATL): the anterior division of the inferior temporal and fusiform gyri
4. Posterior middle temporal gyrus (pMTG): the temporo-occipital part of the middle temporal gyrus

The final ROI covered the angular gyrus (AG) and included voxels with a >30% probability of falling within this region in the LPBA40 atlas (Shattuck et al., 2008). A different atlas was used in this case because the AG region defined in the Harvard-Oxford atlas is small and does not include parts of the inferior parietal cortex typically implicated in semantic processing. The 30% inclusion threshold we used to define ROIs is consistent with our previous work (Hoffman, 2019; Hoffman & Tamm, 2020).

### Data and code availability statement

The decoding accuracy maps for each searchlight analysis are available at https://osf.io/2ueh3/. However, the conditions of our ethical approval do not permit public archiving of data at the level of individual participants because participants did not provide sufficient consent for this.

## Results

### Behavioural performance

We first checked the accuracy of the meaningfulness judgements in each imaging run to ensure that participants were responding attentively. We identified five runs from three participants that contained correct responses on fewer than 60% of trials. Participants failed to respond on a large number of trials in these runs, so we excluded these runs from analysis. One additional run was excluded because it was terminated early due to a scanner fault. Mean accuracy was >78% on all other runs and all participants had at least three valid runs for analysis.

Participants were highly accurate in responding to event sentences (*M* = 95.5%; *SD* = 4.1%) and to anomalous sentences (*M* = 91.3%; *SD* = 6.1%), though more accurate for the event sentences (*t*(25) = 3.02, *p* = 0.006). They were slower to respond to anomalous sentences (*M* = 1940ms; *SD* = 390ms) than to event sentences (*M* = 1811ms; *SD* = 388ms; *t*(25) = 5.19, *p* < 0.001). We also compared performance for event sentences that used an active vs. passive syntactic structure. Participants were faster to respond to active sentences (active *M =* 1689ms, *SD* = 380ms; passive *M =* 1936ms, *SD* = 402ms; *t*(25) = 12.0, *p* < 0.001). There was no significant effect of syntactic structure on accuracy (active *M =* 96.1%, *SD* = 4.4%; passive *M =* 95.0%, *SD* = 4.4%; *t*(25) = 1.94, *p* = 0.064).

### Univariate activation

Results of univariate analyses are shown in Supplementary Figure 1. Relative to rest, event sentences and anomalous sentences activated similar networks, including posterior temporal and occipitotemporal cortices and frontoparietal cognitive control networks. When event sentences were contrasted with the more difficult anomalous sentences, the anomalous sentences produced more activation in a left-lateralised network of regions associated with semantic processing. This included parts of IFG, pMTG, ATL and AG. This suggests that semantic judgements about anomalous sentences required greater engagement of the semantic system, consistent with longer reaction times on these trials. Greater activation for event sentences relative to anomalous sentences was found in ventromedial prefrontal and posterior cingulate cortices, consistent with previous data implicating these regions specifically in the processing of coherent events (Lerner, Honey, Silbert, & Hasson, 2011). Alternatively, this effect might be a consequence of greater disengagement of these default mode regions on the more difficult anomalous trials (Mckiernan, Kaufman, Kucera-Thompson, & Binder, 2003).

### Searchlight analyses

Analysis 1 tested whether activity patterns could discriminate between specific sentences describing the four events, without requiring generalisation to new descriptions of the events. Figure 1B shows decoding accuracies in areas where decoding significantly exceeded chance performance. Above-chance decoding was observed across the cortex and bilaterally, with the highest accuracies found in primary and early visual cortices. Other areas showing high decoding accuracy included left pMTG, regions along the length of the superior temporal gyrus, the lateral prefrontal cortex, particularly IFG, and the dorsomedial frontal lobe, close to the pre-supplementary motor area. In the right hemisphere, decoding performance was generally poorer but exceeded chance levels in many areas. The Neurosynth word cloud (Figure 1C) indicates that regions displaying high decoding accuracies were associated with terms relating to visual processing and also to language and semantic processes. Supplementary Figure 2 splits the results of this analysis into classifiers trained with active vs. passive sentences. Significant decoding was found in similar regions for both sentence types, though decoding was generally more successful when the analysis was restricted to passive sentences. This might be indicative of stronger neural signals elicited on these trials, which participants took around 250ms longer to process. In summary, many regions of the brain showed distinct patterns of activation for specific sentences that described different events. However, this analysis does not allow us to determine whether these effects are driven by differences in the conceptual content of the sentences or by differences in their surface orthographic, phonological, lexical and syntactic properties.

In Analyses 2 and 3, we trained classifiers on one set of sentences and tested their ability to classify a new set of sentences with different lexical or syntactic forms. Areas of above-chance decoding for each analysis are shown in Figure 2B and 2D; the conjunction map in Figure 2E shows regions that were significantly above-chance in both analyses (for peak activation co-ordinates, see Supplementary Materials). Both analyses identified a similar set of brain regions. Strongest decoding was observed in left AG and pMTG in both cases, with additional clusters present in the left ATL and IFG. Decoding was also successful in the right-hemisphere homologues of most of these regions. The posterior cingulate and ventromedial prefrontal cortices also showed evidence of event classification that generalised across stimuli. Compared with Analysis 1, decoding in early visual regions appeared to be much weaker. To confirm this, we performed a post-hoc exploratory analysis of decoding in early visual cortex (see Supplementary Figure 3). Decoding of events in early visual cortex was significantly poorer in Analyses 2 and 3 compared with Analysis 1, though it exceeded chance level in Analysis 2. As we will discuss later, we believe that effects in this region are a consequence of differences in the perceptual properties of different sentence stimuli.

Figure 2F indicates that the Neurosynth terms most associated with accuracy in Analyses 2 and 3 refer to semantic processing, theory of mind and the default mode network but not to perception. Thus, when required to generalise event information to novel sentences with different structure and form, decoding in early perceptual regions was much reduced while it remained robust in higher-level semantic and default mode regions.

The final two analyses, shown in Figure 3, tested for regions whose activity discriminated between different syntactic structures and lexical items, rather than between different events. In Analysis 4, we investigated which regions were able to discriminate between active and passive sentence structures. High levels of decoding were observed in early visual regions, with similar levels of accuracy to that observed in Analysis 1 (see Supplementary Figure 3). Decoding also exceeded chance levels in the intraparietal sulcus bilaterally, left posterior IFG (BA44) and the left precentral gyrus. These regions show little overlap with the event-coding regions found in Analyses 2 and 3. Finally, Analysis 5 tested whether any regions could discriminate between sentences that used different lexical items to describe the same event. Above-chance decoding in this analysis was only found in occipital cortices, extending into left occipitotemporal cortex.

### Regions of interest

Decoding performance in anatomical ROIs is shown in Figure 4B. When the classifiers were trained to discriminate between events (Analyses 1 to 3), all ROIs showed better performance than expected by chance (with the exception of vATL, which was not above chance in Analysis 1). In contrast, no regions performed better than chance when decoding syntactic structure of the sentences, independent of their content (Analysis 4). In Analysis 5, which tested ability to decode the different lexical items used to describe an event, only pMTG showed classification significantly better than chance (though power to detect decoding effects was lower in this analysis because fewer trials were used in each iteration of training). Thus the general pattern is that regions within the semantic network encode the conceptual content of sentences but not their surface characteristics.

For each ROI, we directly compared decoding accuracy between analyses on a pairwise basis. For each pairwise comparison, we used the Stelzer et al. (2013) permutation method to generate a distribution of differences under the null hypothesis, and the position of the observed difference in this distribution was used to determine its p-value. The results of these pairwise comparisons were FDR-corrected for multiple comparisons and significant differences are shown in Figure 4C. All regions except IFG showed some significant differences between analyses. In three regions – pMTG, aSTG and AG – the syntax classifier (A4) performed more poorly than one or more event-based classifiers (A1-A3). This is consistent with the findings of the searchlight analyses, which showed that patterns in the canonical semantic network were able to decode event identity but not syntactic structure. There were also differences between the event-based classifiers. Classification in pMTG was more successful in Analysis 1, where trained and tested sentences were the same, compared with Analyses 2 and 3, where generalisation to novel sentences was required. Interestingly, the reverse was true for vATL and AG: these regions showed better classification when required to generalise to new sentence forms.

## Discussion

Previous studies have found that a network of left-lateralised semantic processing regions represent word and object concepts in a way that generalises across diverse stimulus forms. Here we investigated whether such stimulus independence is also a feature of the neural coding of event semantics. We trained a classifier to discriminate between sentences describing four distinct events and tested its ability to decode the same events from new sentences with different syntactic structure or lexical constituents. Neural patterns in a network of semantic processing regions, predominately in the left hemisphere, showed event coding that generalised over both syntactic and lexical changes to the stimuli. In contrast, early visual regions were able to decode specific event sentences but could not generalise to new sentences. Decoding of syntactic structure and lexical items, independent of event content, was limited and occurred primarily outside the canonical semantic network. Our results indicate that a distributed network of left-dominant regions code information about the conceptual significance of event sentences in a way that is robust to changes in their surface forms.

Searchlight analyses revealed strongest evidence for stimulus-independent event representations in left pMTG and AG. In addition to being considered core parts of the semantic network, both of these regions have been specifically implicated in processing of event knowledge. pMTG shows increased engagement for processing verbs relative to nouns and is also involved in the neural representation of tools and actions (Bedny, Caramazza, Grossman, Pascual-Leone, & Saxe, 2008; Caspers, Zilles, Laird, & Eickhoff, 2010; Ishibashi, Pobric, Saito, & Lambon Ralph, 2016; Peelen, Romagno, & Caramazza, 2012). AG, in addition, has been implicated in the processing of meaningful stimuli that are extended in time, such as sentences, narratives and action sequences (Branzi, Humphreys, Hoffman, & Ralph, 2020; Humphreys & Lambon Ralph, 2017; Humphries, Binder, Medler, & Liebenthal, 2006; Lerner et al., 2011). This has led to claims that this region is essential for combinatorial semantic processes, in which individual word meanings are integrated into a coherent global representation (Bemis & Pylkkänen, 2012; Humphries et al., 2006; Price, Bonner, Peelle, & Grossman, 2015). Beyond these two core event processing regions, however, a more extended network of areas were able to decode events in a cross-sentence fashion, including portions of the ATLs, left IFG and parts of the posterior cingulate and ventromedial prefrontal cortex. All of these areas have been identified as belonging to the brain’s core semantic network (Binder & Desai, 2011; Binder, Desai, Graves, & Conant, 2009; Lambon Ralph et al., 2017). In addition to overlapping considerably with the default mode network, these areas tend to be the most physically distant and functionally distinct from regions serving primary sensory and motor processing (Margulies et al., 2016). This suggests that they are particularly well-suited to serving supramodal and internally-directed cognitive processes that abstract away from perceptual input. Our results support this general view and indicate that activation patterns throughout this network can distinguish between different conceptual states in a stimulus-independent manner.

According to an embodied semantics perspective, regions that discriminate between events may also be coding sensory-motor simulations elicited during processing. For example, visual imagery elicited by the sentence “The cow jumped over the fence” will be very different to “The student considered the problem”. One candidate region for such effects is early visual cortex (V1 and V2), which showed strong decoding in some of our analyses. Borghensani et al. (2016) investigated perceptual effects in this area when participants processed object words. They found that neural patterns were predicted by the perceptual properties of the stimulus word (number of letters) as well as by the visual characteristics of its referent (the real-world size of the object). This result suggests that visual cortex is involved in representing the visual properties of activated concepts, as well as direct perception of external stimuli (see also Pearson, Naselaris, Holmes, & Kosslyn, 2015). In our study, however, we believe effects in early visual cortex are driven primarily by the properties of the stimuli themselves. The strongest decoding in this region was observed in Analyses 1, 4 and 5. In all of these cases, there were systematic differences in the number of letters in the sentences that could aid classification (either because the classes were composed of different lexical items or because they used active vs. passive constructions). In contrast, in the key analyses of stimulus-independent event decoding (2 and 3), sentence length was not informative about class and decoding in early visual cortex was much poorer. Nevertheless, we did observe above-chance decoding of event identity that generalised over lexical items (Analysis 2) and which cannot be ascribed to stimulus properties. This might indicate that visual cortex was also involved in representing that visual qualities of the events being described. This is important question for future research to address.

More generally, the network of regions observed in the present study converges with other studies applying multivariate decoding techniques to sentences (Anderson et al., 2017; Frankland & Greene, 2015; Pereira et al., 2018) and to language at the discourse level (Huth, De Heer, Griffiths, Theunissen, & Gallant, 2016; Wehbe et al., 2014). These also report successful decoding of sentence meaning in left inferior parietal, lateral temporal and inferior prefrontal voxels. Like previous studies, we also found that the network coding meanings was left-lateralised, though there was evidence for above-chance decoding in right-hemisphere homologues as well. However, the present study diverges from some previous work in identifying in left ventral ATL (inferior temporal and fusiform gyri) as an additional area involved in the coding of sentence meaning. This area often suffers from susceptibility artefacts and associated signal dropout, which our use of multi-echo fMRI may have alleviated (Kundu et al., 2017; Visser, Jefferies, & Lambon Ralph, 2010).

The ventral ATL is a key site for semantic representation and, along with the adjacent perirhinal cortex, is thought to be involved in representing conceptual relationships amongst objects and words (Bruffaerts et al., 2019; Clarke & Tyler, 2015; Lambon Ralph et al., 2017; Rogers et al., 2004). Accounts have emphasised the role of this region in understanding single words and objects but have not proposed a specific role in the representation of events. Indeed, some researchers propose a division of labour within the semantic system whereby the ATL codes taxonomic semantic structure while the temporoparietal cortex is represents thematic associations between concepts (Mirman, Landrigan, & Britt, 2017; Schwartz et al., 2011). On this view, ATL should primarily be involved in coding the properties of individual objects or words while the temporoparietal cortex plays a greater role in coding the semantics of events, due to its sensitivity to thematic information. Our data are agnostic on this issue. We found that patterns in ventral ATL do discriminate between different event descriptions. However, the events we used contained different agents and patients and it is possible that ventral ATL voxels coded the properties of these constituents (e.g., discriminating between cows and lorries), rather an integrated event-level representation. Future studies could disentangle these possibilities by using sentences that recombine the same constituents into different event structures (e.g., Frankland & Greene, 2015; Frankland & Greene, 2020).

A functional division within the semantic system has also been proposed between representational regions that code meaning and control regions that regulate and shape semantic activation based on current goals (Badre & Wagner, 2007; Hoffman et al., 2018; Jefferies, 2013). Following this distinction, the Controlled Semantic Cognition theory proposes that IFG and pMTG are involved in semantic control processes rather than representation (Lambon Ralph et al., 2017). In the present study, we observed robust decoding of events in both of these areas, as have other semantic decoding studies at the word and sentence levels (Anderson et al., 2017; Devereux et al., 2013; Fairhall & Caramazza, 2013; Pereira et al., 2018). If these regions are important for regulating semantic activation rather than storing conceptual representations, why can their neural patterns reliably distinguish between different event concepts? There are a number of possible answers to this. Different events may vary, for example, in the demands they place on control processes and this systematic variation in engagement might be sufficient to allow classification. Alternatively, different events might require different control processes with distinct neural signatures. Better understanding of this issue will require more precise formulation of the mechanisms and computations involved in semantic control. Many computational models of semantic *representation* exist and these make specific predictions about the representational structures involved (e.g., O’Connor, Cree, & McRae, 2009; Rogers et al., 2004; Schapiro, McClelland, Welbourne, Rogers, & Lambon Ralph, 2013; Taylor, Devereux, Acres, Randall, & Tyler, 2012). Conversely, there has been little formal modelling of semantic *control* processes (though for recent exceptions, see Hoffman et al., 2018; Jackson, Rogers, & Lambon Ralph, 2021). The lack of established theoretical models makes it difficult to make precise predictions about how neural patterns in control regions should vary and how these areas might be distinguished from those that represent knowledge. This remains a key challenge for future work to address.

While we found that brain regions throughout the canonical semantic network were able to decode event semantics independent of syntactic structure, coding of syntactic information was much more limited. None of our semantic ROIs were able to discriminate between active and passive sentences in an event-independent fashion; instead, this form of decoding was mostly limited to occipital cortex and superior frontal and parietal regions. Some of these effects may reflect differences in non-semantic processing demands to the two sentence types. Passive sentences are necessarily longer than their active equivalents. This means they require greater visual analysis, additional eye movements and greater working memory recruitment. The decoding we observed may reflect variation in these pre conceptual demands. In addition, English speakers typically expect the initial noun phrase in a sentence to represent the agent (agent-first bias; see Kamide, Scheepers, & Altmann, 2003). When reading passive sentences, this initial thematic role assignment is incorrect and has to be revised. This reanalysis process has been proposed as an explanation for greater left IFG activation for passive sentences (Feng et al., 2015; Mack, Meltzer-Asscher, Barbieri, & Thompson, 2013). In our study, we observed decoding in a posterior portion of left IFG (pars opericularis; BA44), which may reflect these processes of reanalysis. The region was somewhat posterior to the peak decoding in the event-discrimination analyses. This is consistent with the idea of graded specialisation in left IFG, with semantic processing engaging more anterior regions and syntactic processing more posterior areas (Friederici, 2012; Rodd, Vitello, Woollams, & Adank, 2015). In general, however, our results support the view that few regions in the language network are primarily influenced by syntactic information, independently of the semantic content of language (Blank, Balewski, Mahowald, & Fedorenko, 2016; Fedorenko, Nieto-Castanon, & Kanwisher, 2012; Mollica et al., 2020).

Finally, we tested for areas that discriminated between different lexical items used to describe the same events. The searchlight analysis found that this decoding was restricted to early visual and left occipitotemporal regions, which is likely to be a consequence of orthographic rather than semantic processing (e.g., differences in word length). In the ROI analyses, however, pMTG showed above-chance discrimination. This effect could indicate differences in lexical access processes supported by posterior temporal regions (Davis, 2016; Lau, Phillips, & Poeppel, 2008; Lewis & Poeppel, 2014). Alternatively, the distinct neural patterns observed here may be a consequence of subtle semantic distinctions between the different lexical items. Although we designed the materials such that lexical substitutes were as conceptually similar as possible, there were inevitably some differences between them (cows and bulls are not identical, nor are fences and gates, and so on). With a larger sample, we may have been able to decode these subtle semantic distinctions more widely within the semantic network.

One limitation of the study is that our decoding is based on a small number of simple events, all with a similar agent-verb-patient structure. The human semantic system is capable of representing a far more complex and varied selection of events than those used here. Further research will be needed to understand how and where specialisation occurs for particular event classes. For example, events could be selected based on the degree to which they engage different types of sensory-motor simulations, allowing for targeted investigation of the predictions of embodied semantics theories (Barsalou, 1999; Pulvermüller, 2013; Zwaan, 2004). On the whole, however, the present study suggests that a distributed network of language and semantic regions code event-level semantic information in a manner that generalises across lexical and syntactic variation in the sentences used to describe them. This suggests that the core function of this network is to abstract away from perceptual input and represent the underlying conceptual significance of our experiences.

## Acknowledgements

PH was supported by a BBSRC grant (BB/T004444/1). Imaging was carried out at the Edinburgh Imaging Facility (www.ed.ac.uk/edinburgh-imaging), University of Edinburgh, which is part of the SINAPSE collaboration (www.sinapse.ac.uk).

