## Supplementary Materials for "Stimulus-independent neural coding of event semantics: Evidence from cross-sentence fMRI decoding"

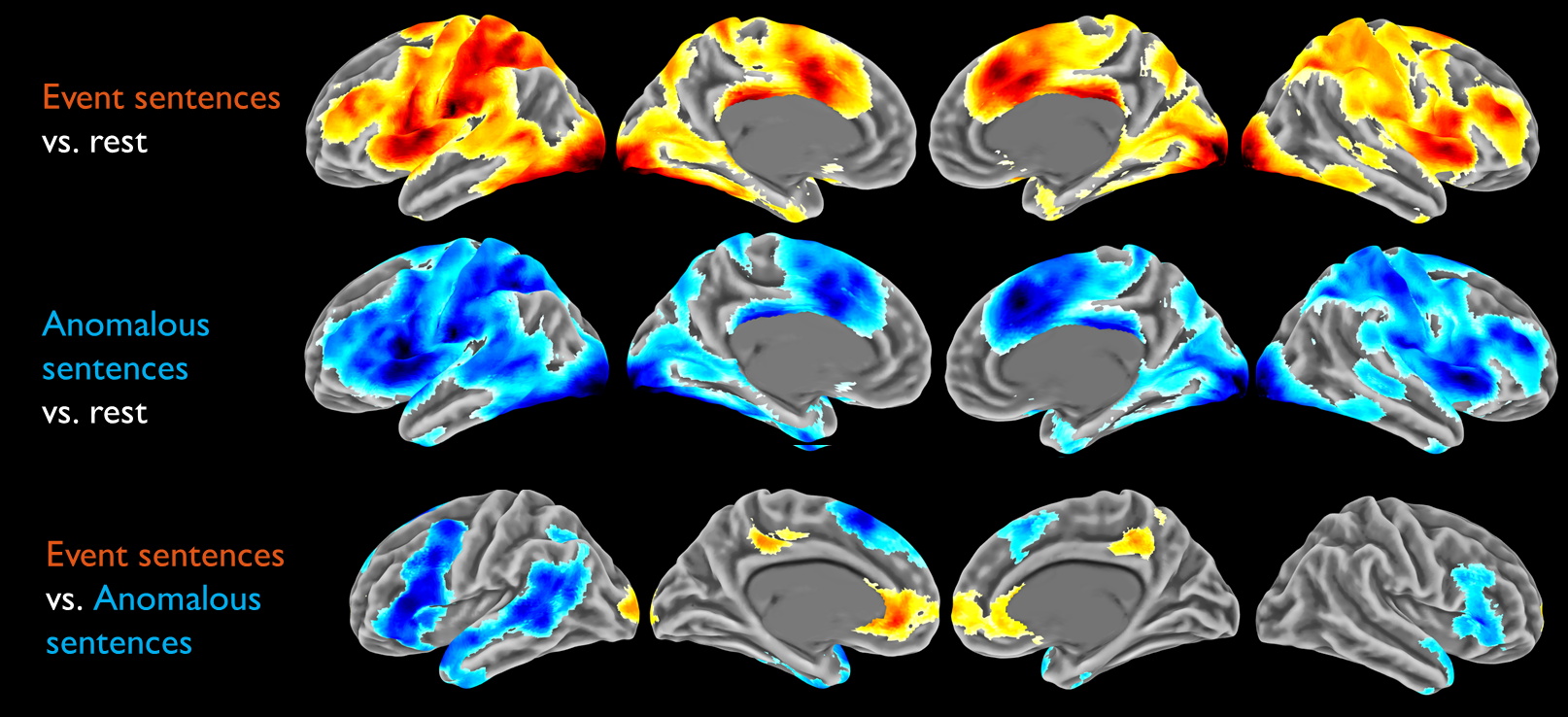

*Supplementary Figure 1: Univariate activation contrasts Images are thresholded at voxelwise p < 0.005 with correction for multiple comparisons at the cluster level using SPM’s random field theory (FWE p < 0.05).*

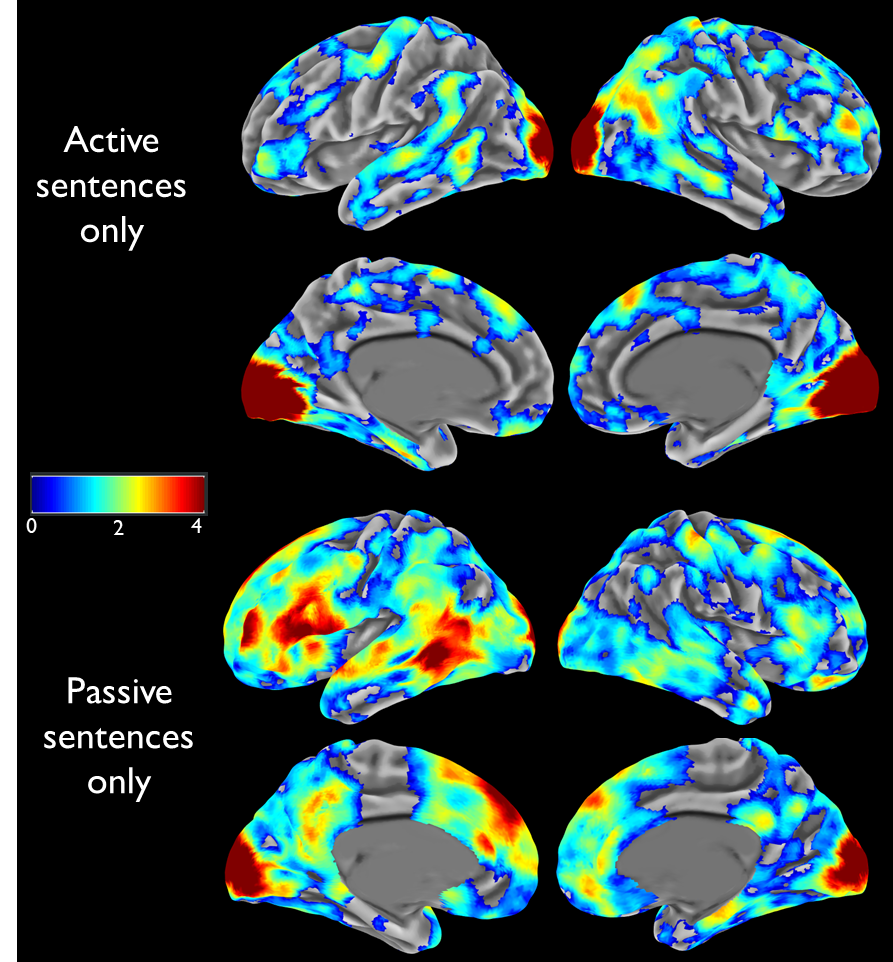

*Supplementary Figure 2: Additional decoding maps for Analysis 1, splitting by syntactic form Decoding accuracy is presented relative to chance level and thresholded at cluster-corrected p< 0.05. Maps were smoothed at 5mm FWHM for display purposes.*

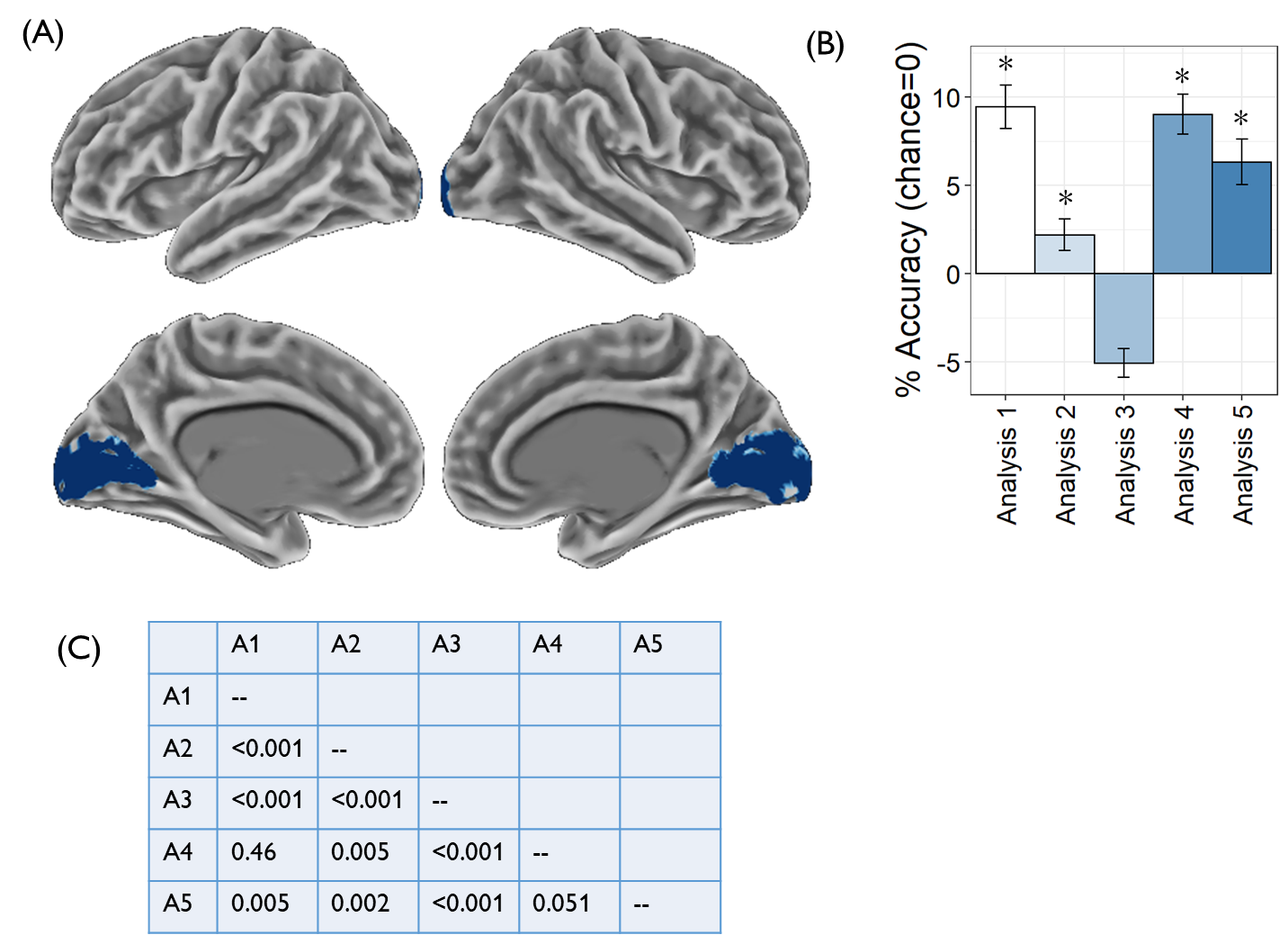

*Supplementary Figure 3: Decoding accuracy in early visual areas*

(A) We performed an exploratory ROI analysis of classification performance in early visual cortex, defined as regions hOC1 and hOC2 (V1 and V2) from the SPM Anatomy toolbox (Amunts, Malikovic, Mohlberg, Schormann, & Zilles, 2000). (B) Decoding accuracy for each analysis; * indicates performance exceeded chance at FDR-corrected *p <* 0.05. (C) Results of pairwise comparisons of decoding performance across analyses. Table shows FDR-corrected p-values when permutation testing for difference between two analyses. A1 = Analysis 1 etc.

Amunts, K., Malikovic, A., Mohlberg, H., Schormann, T., & Zilles, K. (2000). Brodmann's areas 17 and 18 brought into stereotaxic space—where and how variable? *Neuroimage, 11*(1), 66-84.

*Supplementary Table 1: Sentences used as stimuli*

| *Sentence* | *Type* | *Event #* |
| --- | --- | --- |
| The computer processed the file | Meaningful | 1 |
| The laptop analysed the document | Meaningful | 1 |
| The file was processed by the computer | Meaningful | 1 |
| The document was analysed by the laptop | Meaningful | 1 |
| The bull leapt over the gate | Meaningful | 2 |
| The cow jumped over the fence | Meaningful | 2 |
| The gate was leapt over by the bull | Meaningful | 2 |
| The fence was jumped over by the cow | Meaningful | 2 |
| The student considered the problem | Meaningful | 3 |
| The pupil pondered the issue | Meaningful | 3 |
| The problem was considered by the student | Meaningful | 3 |
| The issue was pondered by the pupil | Meaningful | 3 |
| The lorry bumped the lamp post | Meaningful | 4 |
| The truck hit the street light | Meaningful | 4 |
| The lamp post was bumped by the lorry | Meaningful | 4 |
| The street light was hit by the truck | Meaningful | 4 |
| The problem hit the cow | Anomalous |  |
| The issue bumped the bull | Anomalous |  |
| The laptop leapt over the streetlight | Anomalous |  |
| The gate pondered the pupil | Anomalous |  |
| The computer jumped over the streetlight | Anomalous |  |
| The files pondered the truck | Anomalous |  |
| The fence processed the student | Anomalous |  |
| The document was considered by the lorry | Anomalous |  |
| The problem was hit by the cow | Anomalous |  |
| The streetlight was jumped over by the computer | Anomalous |  |
| The fence was analysed by the cow | Anomalous |  |
| The issue was bumped by the bull | Anomalous |  |
| The gate was pondered by the pupil | Anomalous |  |
| The truck was pondered by the files | Anomalous |  |
| The streetlight was leapt over by the laptop | Anomalous |  |
| The lorry was considered by the document | Anomalous |  |

*Supplementary Table 2: Peak activation co-ordinates for Analysis 2*

|  |  | MNI co-ords | | |  |
| --- | --- | --- | --- | --- | --- |
| Location | Extent | x | y | z | *p* |
| L lingual gyrus | 14389 | -6 | -88 | -6 | 0.0001 |
| L posterior MTG |  | -60 | -49 | -6 | 0.0001 |
| L angular gyrus |  | -45 | -70 | 24 | 0.0001 |
| L superior frontal gyrus |  | -12 | 17 | 57 | 0.0001 |
| L IFG (pars triangularis) |  | -51 | 23 | 12 | 0.0001 |
| R angular gyrus |  | 48 | -49 | 30 | 0.0001 |
| R fusiform gyrus |  | 33 | -73 | -24 | 0.0001 |
| L IFG (pars orbitalis) |  | -51 | 32 | -3 | 0.0001 |
| R angular gyrus |  | 51 | -52 | 45 | 0.0001 |
| L frontal pole |  | -15 | 62 | 21 | 0.0001 |
| L IFG (pars orbitalis) |  | -39 | 29 | -12 | 0.0001 |
| R anterior IFG |  | 54 | -13 | -33 | 0.0001 |
| L supramarginal gyrus |  | -51 | -40 | 48 | 0.0001 |
| R superior frontal gyrus |  | 6 | 17 | 57 | 0.0001 |
| R parahippocampal gyrus |  | 33 | -43 | -9 | 0.0001 |
| Precuneus |  | -3 | -58 | 21 | 0.0001 |
| R lateral occipital cortex |  | 30 | -64 | 30 | 0.0001 |
| L anterior MTG |  | -63 | -7 | -18 | 0.0001 |
| R cerebellum |  | 27 | -85 | -36 | 0.0001 |
| R anterior ITG |  | 48 | 2 | -36 | 0.0001 |
| Posterior cingulate | 70 | -3 | -28 | 42 | 0.0047 |
| L precentral gyrus | 26 | -51 | -4 | 33 | 0.0132 |
| L superior parietal lobule | 38 | -21 | -49 | 69 | 0.0138 |

*MTG = middle temporal gyrus; ITG = inferior temporal gyrus; IFG = inferior frontal gyrus.*

*Supplementary Table 3: Peak activation co-ordinates for Analysis 3*

|  |  | MNI co-ords | | |  |
| --- | --- | --- | --- | --- | --- |
| Location | Extent | x | y | z | *p* |
| L angular gyrus | 2394 | -42 | -73 | 27 | 0.0001 |
| L posterior MTG |  | -54 | -61 | 0 | 0.0001 |
| L fusiform gyrus |  | -24 | -43 | -21 | 0.0002 |
| L postcentral gyrus |  | -36 | -28 | 45 | 0.0004 |
| L intraparietal sulcus |  | -45 | -37 | 54 | 0.0008 |
| L IFG (pars triangularis) | 3191 | -48 | 35 | 3 | 0.0001 |
| L frontal pole |  | -18 | 59 | 21 | 0.0001 |
| R frontal pole |  | 12 | 62 | -12 | 0.0001 |
| L superior frontal gyrus |  | -18 | 14 | 54 | 0.0002 |
| R middle frontal gyrus |  | 33 | 29 | 33 | 0.0003 |
| L frontal pole |  | -21 | 59 | 3 | 0.0003 |
| Medial frontal cortex |  | -3 | 47 | 9 | 0.0005 |
| L presupplementary motor area |  | -6 | 41 | 48 | 0.0005 |
| L middle frontal gyrus |  | -36 | 38 | 36 | 0.0009 |
| L dorsomedial frontal cortex |  | -18 | 50 | 36 | 0.0011 |
| Precuneus | 1794 | -3 | -43 | 45 | 0.0001 |
| Precuneus |  | -12 | -52 | 60 | 0.0002 |
| R angular gyrus |  | 54 | -58 | 33 | 0.0003 |
| R lateral occipital cortex |  | 42 | -73 | 27 | 0.0006 |
| R temporoparietal junction |  | 63 | -46 | 24 | 0.0007 |
| R supramarginal gyrus |  | 57 | -46 | 45 | 0.0007 |
| R precuneus |  | 12 | -67 | 27 | 0.0014 |
| Posterior cingulate |  | -6 | -43 | 24 | 0.0020 |
| L anterior STG | 177 | -57 | -7 | -6 | 0.0011 |
| R cerebellum | 284 | 36 | -61 | -45 | 0.0014 |
| R cerebellum |  | 15 | -49 | -45 | 0.0059 |
| R cerebellum |  | 3 | -37 | -54 | 0.0168 |
| R IFG (pars orbitalis) | 172 | 48 | 41 | -3 | 0.0017 |
| R frontal operculum |  | 36 | 26 | 6 | 0.0081 |
| L anterior fusiform gyrus | 119 | -39 | -13 | -36 | 0.0023 |
| R superior frontal gyrus | 132 | 15 | 26 | 57 | 0.0030 |
| R middle frontal gyrus |  | 33 | 17 | 54 | 0.0112 |
| R mid MTG | 328 | 69 | -31 | -12 | 0.0033 |
| R anterior MTG |  | 54 | -7 | -24 | 0.0059 |
| R anterior ITG |  | 42 | -4 | -36 | 0.0115 |
| L cerebellum | 200 | -12 | -70 | -45 | 0.0034 |
| L cerebellum |  | -30 | -79 | -45 | 0.0116 |
| R IFG (pars opercularis) | 88 | 57 | 11 | 18 | 0.0038 |
| R superior frontal gyrus | 49 | 21 | -4 | 54 | 0.0062 |
| L lingual gyrus | 29 | -24 | -61 | 0 | 0.0142 |
| R cerebellum | 30 | 12 | -73 | -39 | 0.0156 |
| L precentral gyrus | 23 | -39 | -16 | 60 | 0.0197 |

*STG = superior temporal gyrus. MTG = middle temporal gyrus; ITG = inferior temporal gyrus; IFG = inferior frontal gyrus.*
